# Spontaneous emergence of behaviorally relevant motifs in human motor cortex

**DOI:** 10.1101/2020.10.25.353326

**Authors:** Tomer Livne, DoHyun Kim, Nicholas V. Metcalf, Gordon L. Shulman, Maurizio Corbetta

## Abstract

Spontaneous neural activity has been shown to preserve the inter-regional structure of cortical activity evoked by a task. It is unclear, however, whether patterns of spontaneous activity within a cortical region comprise representations associated with specific behaviors or mental states. The current study investigated the hypothesis that spontaneous neural activity in human motor cortex represents motor responses that commonly occur in daily life. To test this hypothesis 15 healthy participants were scanned in a 3T fMRI scanner while performing four simple hand movements differing by their daily life relevance, and while not performing any specific task (resting-state scans). Using the task data, we identified cortical patterns in a motor ROI corresponding to the different hand movements. These task-defined patterns were compared to spontaneous cortical activity patterns in the same motor ROI. The results indicated a higher similarity of the spontaneous patterns to the most common hand movement than to the least common hand movement. This finding provides the first evidence that spontaneous activity in human cortex forms fine-scale, patterned representations associated with behaviors that frequently occur in daily life.

## Introduction

Previous work has indicated that spontaneous cortical activity, measured in a task free (resting) state, is correlated between brain regions that are jointly recruited in different task and cognitive states (Golland et al., 2008; Greicius et al., 2003; Nir et al., 2006; Power et al., 2010; Raichle et al., 2001; Xiong et al., 2009). Furthermore, the correlation between cortical regions at rest can be modified by learning, specifically in regions recruited by a novel task pattern (Albert et al., 2009; Harmelech et al., 2013; Lewis et al., 2009; McGregor and Gribble, 2015; Tambini et al., 2010). These findings suggest that ongoing correlated patterns of brain activity represent, at least in part, the global structure of behaviorally relevant brain processing. It is unclear, however, whether ongoing cortical activity reflects only the communication between brain regions, i.e. fluctuations of neuronal assemblies that through time-varying interactions facilitate or inhibit communication (Fries, 2005), or whether ongoing activity patterns also play a representational role, i.e. code or store information about states related to behavior.

Previous behavioral work has suggested that the daily repertoire of human hand movements can be decomposed into principal components (Ingram et al., 2008). Ingram et al. found two components that accounted for the majority of variability of daily motor activity involving the hand. These and other results have suggested a theory of motor function, referred to as motor synergy, which posits that complex movements that serve ecological and behaviorally useful functions are represented in the brain as distinct sets of muscle and joint commands (Latash et al., 2007; Turvey, 2007). Recently, Leo et al. (Leo et al., 2016) demonstrated in a neuroimaging study of human hand movements that activity in the motor cortex can be described in terms of motor synergies. We therefore hypothesized that if patterns of spontaneous cortical activity play a representational role, then those patterns might code motor movements that occur frequently under naturalistic conditions (Ingram et al., 2008). To test this hypothesis, we scanned healthy human subjects (3T fMRI) while they performed a set of simple hand movements that varied in their frequency during daily life (Figure 1a&b), and also while they were not engaged in any task (resting state scans). Resting state scans were conducted immediately before and after task scans. By comparing the similarity between task-induced fMRI BOLD patterns and pre-task resting state patterns we show that resting-state patterns conform more closely to task-evoked states in motor cortex that are associated with more frequently occurring hand movements. These results support our hypothesis that spontaneous cortical activity can represent behaviorally important states.

**Figure 1.**
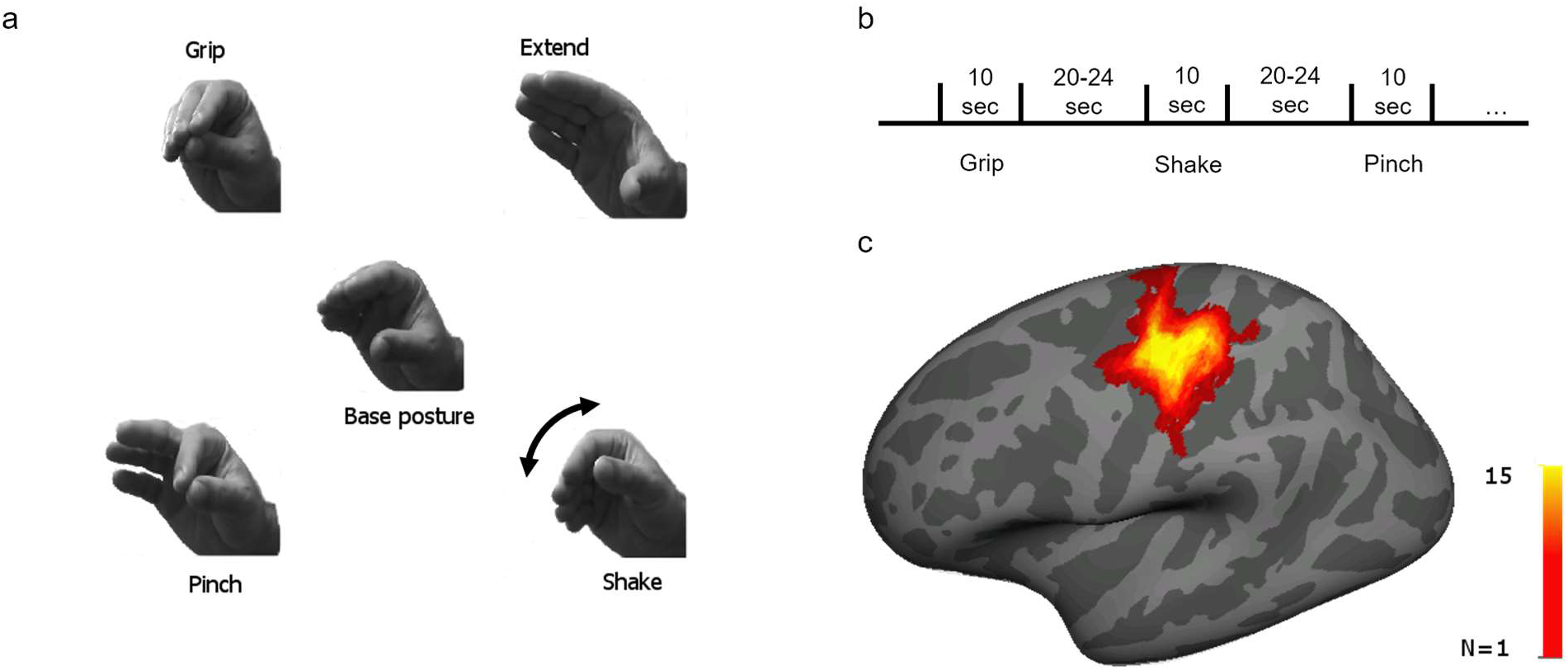
**a)** The four different hand movements used in the study. Center – base hand posture for all movements. In each condition participants were asked to repeatedly and smoothly alternate between the base hand posture and one of the other four positions, creating the different hand movements. Top left – Grip movement, closing hand; top right – Extend movement, opening hand; bottom left – Pinch movement, closing only two fingers; bottom right – Shake movement, rotation of the wrist. **b)** experimental design – participants completed five task runs (except for one participant who completed seven runs, and another who completed only four), and each run included twelve motor blocks (three of each movement) of 10 seconds continuous hand movement. Task blocks were separated by rest blocks (20-24 seconds long). **c)** Motor ROI – for each participant we identified a motor cortex region of interest (ROI) in which the BOLD activation for all the motor tasks was significantly higher than baseline activity. ROIs were identified on the native cortical surface of each participant. For presentation purposes only all ROIs were projected on an average surface (Freesurfer fsaverage) and summed together. The color scale represents the number of participants for which the specific vertex on the surface was included in the ROI. Talairach coordinates of the center of mass of the summed ROIs [−36, −23, 55].

## Materials and Methods

### Task

Participants (N=15, 7 females, mean age 26.5, all right handed) performed a block-design hand-movement task in which they were instructed to move their right hand repeatedly for 10 seconds in one of four movements followed by a variable rest period (20-24 seconds), see Figure 1 a&b.

Four hand movements were used in the current study: Grip, Extend, Pinch, and Shake. Grip and extend were chosen as representing the most common hand movements based on work by Ingram et al (2008) and corresponded to the first two principal components identified in that work. The Shake hand movement in which the participants kept their fingers in a fixed position and rotated their wrist was chosen as a control, less behaviorally relevant hand movement. The Pinch movement was chosen based on its similarity to the Grip movement.

To prevent movement artifacts in the scanner due to the hand movements during task performance the right arm was placed inside a padded plastic half pipe and strapped to it. There were five task runs (except for one participant who performed seven runs and another that performed only four runs), in each run each movement occurred 3 times in a random order. The same participants were also scanned in three five-minutes pre-task resting state scans and three five-minutes post-task resting state scans to evaluate the spontaneous occurrence of the BOLD patterns associated with the different hand movements during rest, and the effect of task performance on the spontaneous activity. During rest scans the participants kept their arm in the restraining mechanism and were instructed to rest their palm on the scanner bed alongside their body. Task performance was verified visually throughout the experiments. Prior to participation in the study all participants signed an informed consent form approved by Washington University IRB. Participants received monetary compensation for their time.

The experiment was controlled using the Psychtoolbox extension to Matlab (Brainard, 1997; Pelli, 1997). Stimuli were presented to the participants using a head mounted mirror set and a back projector.

### MRI acquisition

The data for 2 participants were acquired using a Siemens (Erlingen, Germany) 3-T Tim Trio MR system, using a 16-channel RF head coil. Whole-brain multi-band echo-planar images were acquired using the following parameters: 32 interleaved slices, 2-sec repetition time, 27-msec echo time, 4-mm thickness. Voxel size was 4 × 4 × 4 mm. Up to two high-resolution T1-weighted echo-planar MPRAGE anatomical images (isotropic 1-mmvoxels) were obtained and used to construct a high-resolution image of each participant’s brain.

The data for the rest of the participants were acquired using a Siemens (Erlingen, Germany) 3-T Prisma Fit MR system, using a 32-channel RF head coil. Whole-brain echo-planar images with multiband factor of 3 were acquired using the following parameters: 56 interleaved slices, 1-sec repetition time, 25.8-msec echo time, 3-mm thickness. Voxel size was 3 × 3 × 3. Up to two high-resolution T1-weighted multi-echo MPRAGE anatomical images (isotropic 1-mmvoxels) were obtained and used to construct a high-resolution image of each participant’s brain.

Structural and functional preprocessing were performed using the FreeSurfer and FS-FAST processing stream developed at the Martinos Center for Biomedical Imaging (surfer.nmr.mgh.harvard.edu). Frames were motion-corrected by aligning them to the middle frame in each run. Functional data were registered to the high-resolution structural data and then segmented into left and right cortical surfaces. The intensity level of each frame was normalized. The raw time series was then resampled to the reconstructed surfaces. A 5-mm FWHM Gaussian smoothing was applied to the resampled data. The frames corresponding to the first ten seconds of each scan were discarded from the analysis.

### Analysis

We first defined a motor ROI by performing a glm analysis contrasting motor performance with baseline activity (four movements combined to avoid biasing the ROI selection to a specific movement), using a gamma function defined by the following parameters: Δ= 2.25 and τ = 1.25. Motion correction parameters were used as nuisance regressors in this analysis. We then defined a central-sulcus cluster on the native surface of each participant in which the contrast was significant. To keep the size of the ROIs similar across participants we adjusted the p-level of the contrast until we ended up with an ROI size of 2187 vertices ±7%.

For the main analyses we additionally regressed out the global (mean) signal of the left hemisphere from the temporal activity pattern of each vertex. We then normalized the activity levels in each TR by converting them to z-score values ending up with a relative activation pattern of the different vertices in each TR.

We tested the discriminability of the cortical activation patterns induced by the different hand movements in the ROI using a linear discriminant analysis (LDA) performed in Matlab (Mathworks inc). Classification success was estimated using a cross-validation leave one run out method.

A mean pattern for each hand movement (constructed from all the task runs) was compared to each resting state frame using Pearson correlation. We computed the cumulative distribution function (CDF) of the r^2^ values and identified the r^2^ value that represented the cumulative 90% point. The higher this value is the more the of the activity pattern during rest is explained by the task pattern.

All t-tests reported were two-tailed and paired-sample when appropriate. All ANOVA test comparing effects within subjects where conducted as repeated-measures ANOVA.

## Results

### Movement classification in the motor cortex

To verify that task-evoked BOLD patterns within the motor ROI (Figure 1c) were associated with specific hand movements, we tested whether the different movements could be discriminated using linear discriminant analysis (LDA). Using a leave-run-out cross-validation approach, we calculated classification success for the motor ROI in each acquired brain volume (1 volume per TR) within each block. Classification was significantly better than chance level (see figure 2a; p<10^^-7^, fdr corrected, for the frames starting 10 seconds after trial onset and ending 8 seconds later, chance level 25%). The time-course of the classification success-rate function was highly correlated with the time-course of the mean BOLD signal (Figure 2b; Spearman r^2^=0.77). Since our main aim was to compare task patterns to spontaneously emerging BOLD patterns, we constructed a mean activation pattern for each movement by averaging the patterns during the most activated TRs in a block (the period 10-18 seconds after the beginning of the block). Classification success rates using these mean patterns were high and significantly above chance (t(14)=14.23, p<10^^-7^, mean classification success 0.73, se=0.033 – right most column in Figure 2a). A representational similarity analysis (Figure S1a), in which the patterns for the different movements were spatially correlated, indicated that the Grip, Extend, and Pinch patterns were more similar to each other than to the shake pattern. The high similarity between these three patterns was also apparent in the confusion matrix of the classifier (Figure S1b). Misclassification tended to be between these three movements and the classifiers rarely misclassified one of these three movements as the Shake movement.

**Figure 2.**
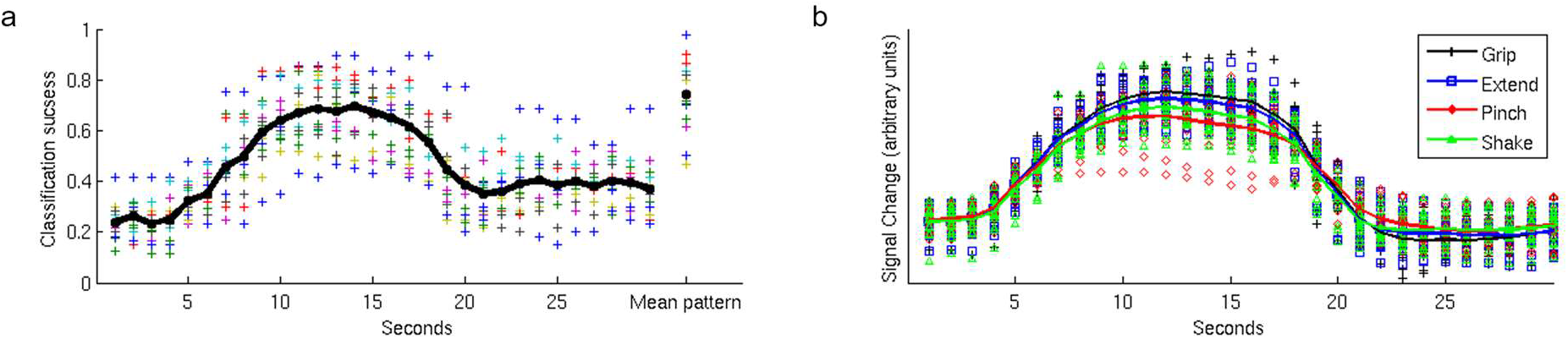
**a)** Classification performance. Using linear discriminant analysis (LDA) we determined whether the pattern of activation in the motor ROI of each participant at each TR discriminated between the different hand movements. The results, presented as a function of time (x-axis), indicate that classification performance followed the hemodynamic response function with a gradual increase in performance starting ~6 seconds after trial onset (spearman r between the mean classification function and the mean BOLD signal representing a full trial, combining all the conditions together, is equal to 0.876). The rightmost data points represent classification performance based on the mean activation patterns constructed by averaging the patterns of the frames starting 10 seconds after trial onset and ending at 18 seconds after trial onset. These mean patterns were used for the analysis described in the following figures. Different colors represent different participants, the black trace represents mean classification levels across participants. Chance level of correct classification was 0.25. For presentation purposes only, each data point of the two participants who were scanned using a 2 second TR was duplicated in the figure (TR #1 is presented as second 1 & 2, etc.). **b)** Mean BOLD time course for the four hand movement conditions. Each data point represents the mean normalized BOLD signal change of each participant to her baseline BOLD activity across all ROI vertices in a specific time point during the block. Different symbols and corresponding colors represent different hand movements. The solid lines represent the mean normalized BOLD signal change across participants corresponding to each hand movement. As can be seen in the figure all four conditions elicited similar BOLD time courses. For presentation purpose only each data point of the two participants who were scanned using a 2 second TR was duplicated in the figure (TR #1 is presented as second 1 & 2, etc.).

### Spontaneous occurrence of movement patterns during rest

To test whether hand movements are represented in spontaneous activity according to their relevance in daily life, the mean task patterns were compared to each pre-task resting state frame using Pearson correlation. An overall similarity measure between the task pattern and spontaneous rest patterns was defined as the 90^th^ percentile value of the cumulative distribution function of the r^2^ values of each pattern (see Figure 3). With this method of quantifying similarity, a higher cutoff value indicates that a larger fraction of rest frames had a higher similarity to the tested pattern than to a pattern having a lower cutoff value. We conducted a 1-way repeated measure ANOVA with pattern as the independent variable, 90^th^ percentile cutoff value as the dependent variable, and participant as a random variable of no concern. The analysis indicate a main effect of movement type (F(3,42)=5.46, p=0.0029). Results are presented in Figure 4a & 4b. Post-hoc paired-sample t-tests indicated that the rest patterns were significantly more similar to the Grip than Shake patterns (t(14)=3.49, p =0.0036,r^2^-cutoff=0.272 and 0.186, respectively), an effect that remained significant after correcting for multiple comparisons. The Extend and Pinch patterns’ cut-off values (0.252 and 0.25, respectively) were also higher than the Shake value but these differences did not survive multiple comparisons (t(14)=2.34, p=0.034, and t(14)=2.35, p=0.034, respectively). These results support our hypothesis that more common hand movements are more frequently represented in the pattern of resting, spontaneous activity in motor cortex.

**Figure 3.**
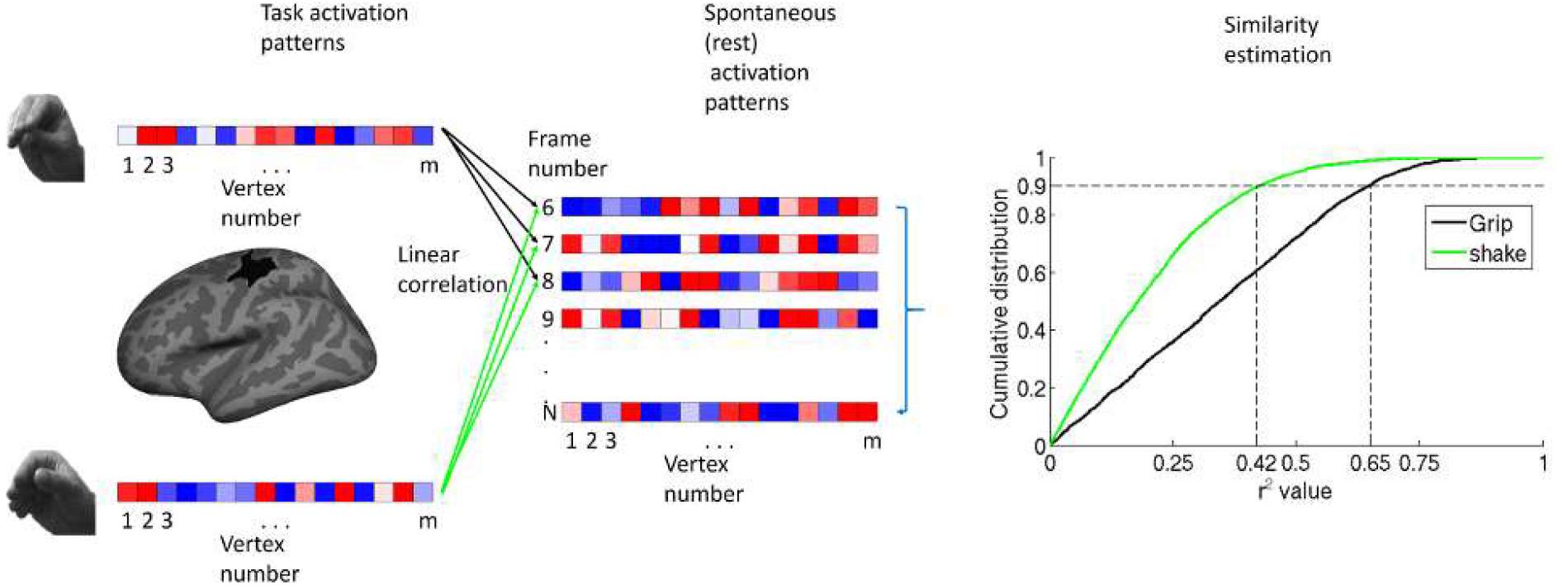
Methods – A spatial pattern was defined for each hand movement (left) in the participant-defined motor ROI (left). Each of the four task patterns were compared to the spatial pattern of the same vertices in each rest frame (excluding the first 10 seconds of each rest scan) using Pearson correlation (middle). For each pattern, we computed the cumulative distribution function (cdf) of r^2^ values across all rest frames (right). In each cdf of each participant we calculated the value corresponding to the 90^th^ percentile of that function (vertical dashed lines). This value was used to estimate the degree of representation of the specific pattern in the rest data. With this method of quantifying similarity, a higher cutoff value indicates that the activity patterns in a larger fraction of rest frames was explained by the tested pattern than by a pattern having a lower cutoff value.

**Figure 4.**
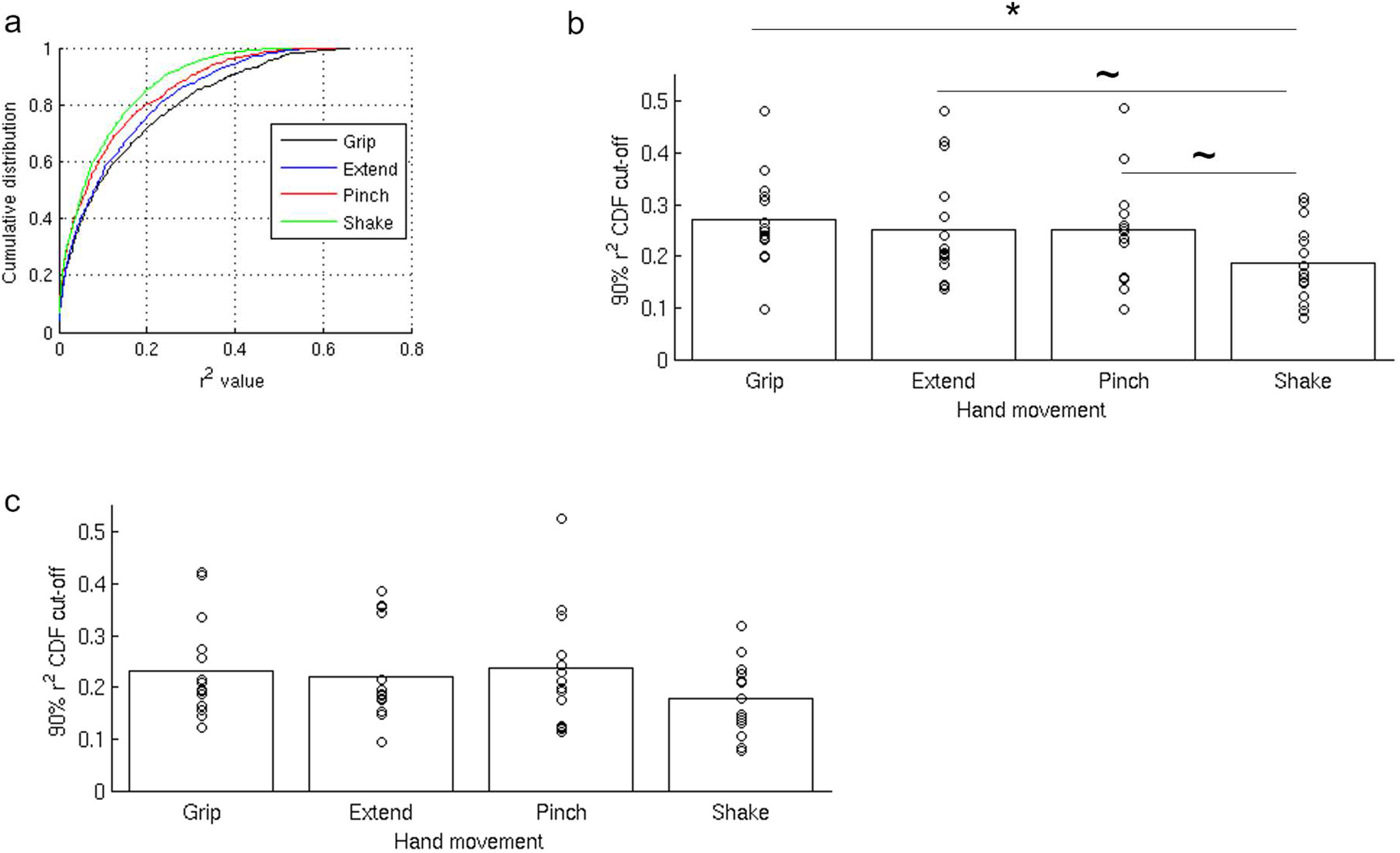
**a)** Representative cdfs (single participant). The Shake condition, corresponding to the green cdf, exhibits the steepest slope, while the Grip condition (black cdf) exhibits the shallowest slope. **b)** r^2^ cutoff values of the four different hand movements’ patterns in the pre-task rest data (all participants). Grip had the highest cutoff value indicating that this pattern was most represented in the pre-task spontaneous activity. The lowest cutoff value was that of the Shake condition, indicating the it was the least represented pattern in the pre-task resting data. The difference between the Grip and the Shake conditions was significant following a correction for multiple comparisons (*). Both comparisons – Extend vs Shake, and Pinch vs Shake were significant before correction, but did not survive the correction for multiple comparisons (~). **c)** r^2^ cutoff values in the post-task rest data. No significant differences were found in the post-task data in terms of r^2^ cutoff values of the different conditions.

### Reduction in spontaneous representation of movement patterns following task performance

When the same analysis was conducted on the post-task rest frames, the main effect of movement type was no longer significant (F(3,42)=2.43, p=0.078, Figure 4c). This null result was due to a reduction in the similarity between the Grip and rest patterns, as well as between the Extend and rest patterns, coupled with the lack of any reduction in the similarity between the Shake and rest patterns (see supplementary analysis and Figure S2).

## Discussion

In the current study, we tested the hypothesis that ongoing cortical spontaneous activity in the human brain represents behavioral states that frequently occur in daily life. In agreement with previous research we found that it is possible to decode subtle hand movements in a motor cortex ROI using fMRI (Bleichner et al., 2014; Di Bono et al., 2015; Ejaz et al., 2015; Leo et al.,2016; Wiestler and Diedrichsen, 2013). Importantly, both common and less-frequent hand movements activated the motor ROI to a similar degree and could be classified accurately in that region during movement execution. We tested whether these movement-evoked patterns were represented to a different degree in spontaneous brain activity according to their occurrence in daily-life (Ingram et al., 2008). The results support our hypothesis, showing a more robust representation at rest in motor cortex of a commonly used vs. infrequently used hand movement. Importantly, this result was found prior to performing the task, following only a brief and equal exposure to the each of the hand movements during the instructions phase. Therefore, it cannot be explained by short-term adaptation or learning following task exposure (see below). These results provide the first evidence in humans that spontaneous brain activity within a region codes for frequent, naturally occurring behavioral states.

While resting patterns of common hand movements were stronger and more frequent (the measure of similarity used combines both variables) than those of a less frequent hand movement, during task performance all patterns, irrespective of familiarity, were accurately classified. This suggests that the motor cortex can represent both novel and common movements, but the latter are those more represented in spontaneous activity. This result is supportive of the notion that patterns at rest form a kind of representational prior from which novel neural patterns can depart, but to which the motor cortex returns at rest. The strategy to keep ‘ready’ on-line common movement patterns is metabolically expensive, but computationally efficient (Attwell and Laughlin, 2001).

When the same analysis was conducted using resting state scans that were collected immediately following task performance, the difference between the conditions was no longer significant. The representation of the hand movement that was most present in the pre-task resting activity was significantly diminished while the other hand movements were relatively unchanged. This selective pattern of changes suggests that the lack of significant effect in the post-task comparison is not due to an overall reduction in neural activity or a reduction of the overall variability of neural activation in the ROI, but instead reflected a long-term (time scale of minutes) pattern-specific adaptation process similar to the one commonly observed online during task performance (Dinstein et al., 2007; Grill-Spector and Malach, 2001). Such long-term adaptation has been previously reported in the resting state following learning to perform a novel visual discrimination task (Guidotti et al., 2015).

### Slow transitions between resting state states

Similar results to those obtained with the mean pattern were obtained using a spatio-temporal pattern in which the pattern across the chosen TRs was concatenated to one another and compared to similarly concatenated resting-state frames (see supplementary material and Figure S3). This result suggests that any non-stationarity (Allen et al., 2014; Handwerker et al., 2012; Hutchison et al., 2013; Liu and Duyn, 2013; Smith et al., 2012; Zalesky et al., 2014) in the fMRI resting-state spontaneously emerging patterns is modulated by slow transitions, which is consistent with the known temporal properties of the hemodynamic response function estimated using task data (Blamire et al., 1992; Boynton et al., 1996; Buckner et al., 1996; Miezin et al., 2000). Alternatively, it might suggest that the spontaneous activity represents a weighted combination of all the represented patterns, in which case the most dominant one across most TRs would be the grip pattern. Further studies will be required to determine which option is correct.

### Cortical activation and motor execution

An open question that will require further research is how the motor cortex dissociates between spontaneous patterns that do not require the initiation of an actual motor act and patterns involving an actual demand for real movement. One possibility is that of gating by a different brain region. Such a mechanism would suggest that every motor plan is composed of at least two components, an instruction for action and instruction for a control area, to allow the motor plan to pass down the efferent pathway. Some support for this suggestion is found by comparing motor execution to motor imagery in healthy participants (Roux et al., 2003) and in amputees (Raffin et al., 2012). These studies reported a difference between motor imagery and motor execution activation in the recruitment of the supplementary motor cortex regions. Another option is that of a threshold based mechanism that requires a high match between the activated pattern and the optimal pattern for that specific motor action. The similarity observed in the current study between the spontaneous and the task-induced patterns might not have been sufficient to activate an afferent command to the muscles to perform the actual movement. Such a mechanism could operate on the basis of a passive, rather than active, gating mechanism.

### Ongoing activity and cognitive control

Previous work has indicated that ongoing spontaneous cortical activity can shape and even be used to predict success in a variety of perceptual and cognitive tasks (Bode et al., 2012; Cole et al., 2016; Coste et al., 2011; Hesselmann et al., 2008; Sadaghiani et al., 2010; Tavor et al., 2016; Wyart and Tallon-Baudry, 2009) as well as account for behavioral deficits in patient populations (Carter et al., 2010; He et al., 2007). The current finding that spontaneous activity represents behaviorally meaningful information can help explain how internal representations that incorporate statistical regularities in the environment and behavior might shape the automatic interpretation and response to events in our daily life.

## Acknowledgements

The work was supported by NIH grant 5R01MH09648204, TL received support from the center for absorption in science, Israeli Ministry of Aliyah and Immigrant Absorption.

## Authors’ contribution

TL, GLS, and MC designed the experiment and wrote the ms; TL performed the analysis; TL, NVM, and DK collected the data.

