## Supplementary material for "Spontaneous emergence of behaviorally relevant motifs in human motor cortex": Figure S

### A representational similarity analysis between the activation patterns of the four movements

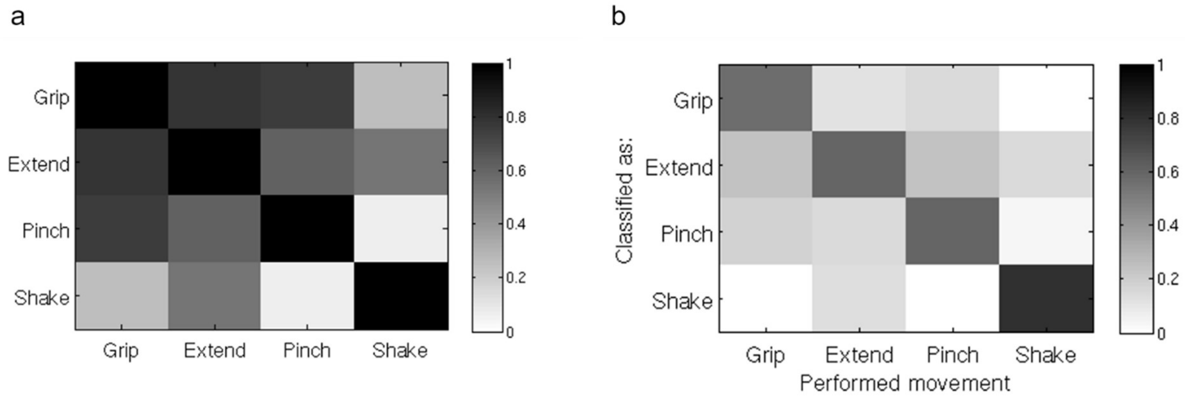


Figure S1. a) Similarity matrix of task-defined patterns. Pearson correlation was used to estimate the similarity between the different patterns used in the main analysis. Similarity between the Grip, Extend, and Pinch patterns was higher than their similarity to the Shake pattern. the color scale represents the mean Pearson r values across all the participants. b) Confusion matrix representing the mean correct classification rate of each movement pattern (diagonal) and the misclassification rate as a function of the incorrect response given by the classifier. The figure shows that the similar patterns (Grip, Extend, and Pinch) were misclassified as each other more often than they were misclassified as the Shake movement. The color scale represents the fraction of cases in which the tested pattern (x-axis – the performed movement) was classified as each of the four movements (y-axis).

**Pre- and post-task comparison**

Paired sample t-tests were used to compare the representation of each condition in the pre- and post-task rest spontaneous activity. The results indicated a significant reduction in representation of the Grip condition ($t(14)=2.79$, $p=0.014$), and the Extend condition ($t(14)=2.42$, $p=0.03$). The extent of the Pinch and the Shake conditions representation was not significantly different between the pre- and post-task rest data sets (both $p>0.3$).

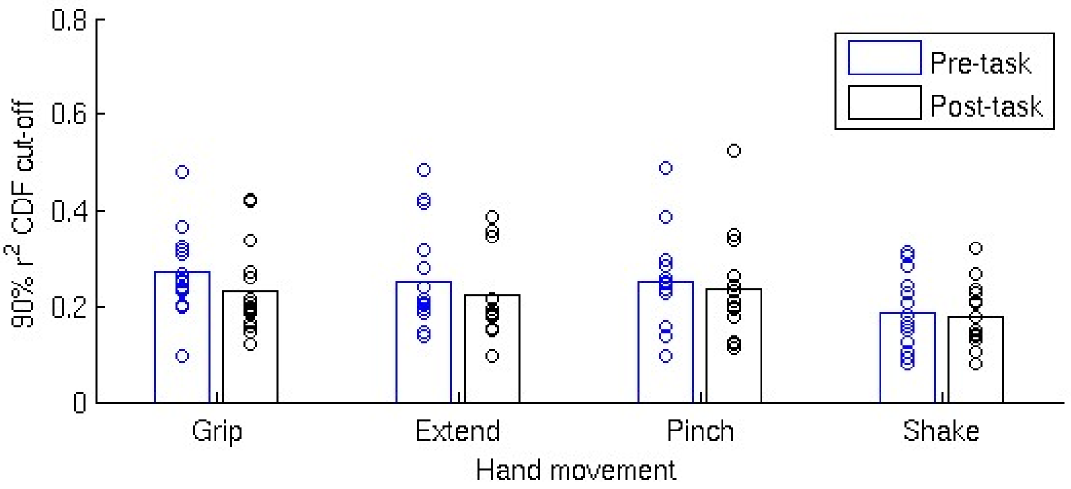


Figure S2. A comparison between the similarity of each task-defined pattern and the pre- and post-task rest spontaneous patterns. In the Grip and the Extend condition there was a significant reduction in the overall observed similarity in the post-task rest relative to the pre-task rest. No such reduction was observed for the Pinch and Shake conditions.

**Spontaneous occurrence of movement patterns during rest estimated using spatio-temporal patterns**

We repeated the main analysis using spatio-temporal patterns, in which the patterns across the chosen TRs were concatenated to one another, creating an eight seconds cortical activation matrix (vertex × TR), and compared to similarly concatenated consecutive resting-state frames using Pearson correlation (each of the two matrices was converted to a 1-dimensional vector for this purpose). As with the mean pattern the analysis of the pre-task rest data indicated a main effect of movement type ($F(3,42)=5.36$, $p=0.0032$). Results are presented in Figure S3a. Post-hoc paired-sample t-tests indicated that the Grip pattern was significantly more similar to the rest patterns than the Shake pattern ($t(14)=3.66$, $p=0.0026$) which was significant after correcting for multiple comparisons. The Extend pattern's cut-off values were also higher than the Shake value but this difference did not survive multiple comparisons ($t(14)=2.48$, $p=0.026$). The Grip and Pinch conditions were also significantly different ($t(14)=2.21$, $p=0.044$), but did not survive correction. When the same analysis was conducted on the post-task rest frames (Figure S3b) the main effect of the movement type was no longer significant ($F(3,42)=1.23$, $p=0.31$). Additionally, as in the main analysis, the significant changes between the pre- and post-task rest similarity with the task patterns resulted from decreased similarity of the post-task rest to the Grip ($t(14)=3.28$, $p=0.0055$) and to the Extend ($t(14)=2.57$, $p=0.022$) conditions relative to the pre-task rest.

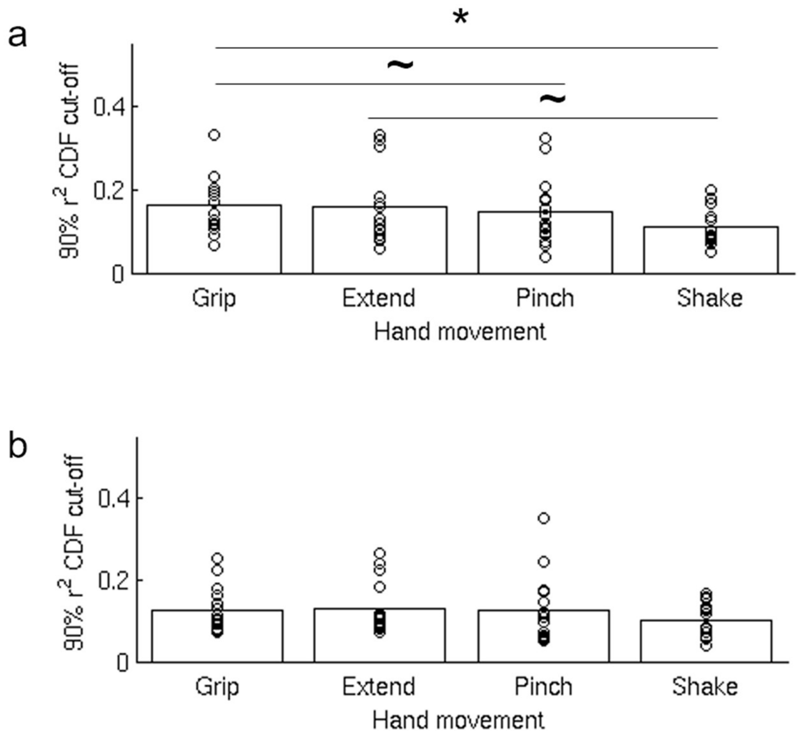


Figure S3. **a)** $r^2$ cutoff values of the four different hand movements' spatio-temporal patterns in the pre-task rest data. There was a significant main effect of movement ($F(3,42)=5.36$, $p=0.0032$) in a repeated measures ANOVA. Grip had the highest cutoff value (0.165) indicating that this pattern was the one most represented in the pre-task spontaneous activity. The lowest cutoff value was that of the Shake condition (0.11), indicating the it was the one least represented in the pre-task data. Only the difference between the Grip and the Shake conditions was significant following a correction for multiple comparisons (* $t(14)=3.66$, $p=0.0026$). The comparisons of Extend (0.159) vs Shake and Grip vs Pinch (0.145) were significant before correction, but did not survive correcting for multiple comparisons (~ $t(14)=2.485$, $p=0.026$, and $t(14)=2.21$, $p=0.044$, respectively). **b)** $r^2$ cutoff values in the post-task rest data using the spatio-temporal patterns. No significant differences were found in the post task data in terms of $r^2$ values of the different conditions.
